# Influence Of Urbanization On Ecological Status Of River In Amhara Region, Ethiopia 2019

**DOI:** 10.1101/2019.12.28.889949

**Authors:** Melsew Setegn Alie

## Abstract

**Introduction:** Urbanization is one of the main causes for environmental problems due to the introduction of pollutants into water bodies. Lahi is crossing fintoselam. This river has long been used for a variety of purposes including source of public water supply, small scale irrigation, bathing, washing, animal watering, sand and stone dredging and recreation.

**Objective:** To assess the influence of Finoteselam town on ecological status of Lahi river

**Methods:** The assessment were assessed using physicochemical parameters, physical habitat assessment, biotic indices /metrics, human activity by observation as well as macroinvertabretes samples from eleven sampling sites coded S1 to S11 along the river using the standard procedures. The assessments were involved in-situ measurements and collection of water samples in April 2019 then, analyzed the water samples in laboratory.

**Results:** The biological analysis reveals a notable reduction of the diversity macroinvertabrates in the downstream direction where the minimum was at midstream sites. Upstream has significantly better macroinvertabrates assemblage than midstream (p-value<0.05). Physical habitat evaluation scores varied from 43 at S8 to 173 at S1 and relatively lower score were recorded at midstream sites. Low % of Ephemeroptera, Plecoptera and Trichoptera with high family biotic index and high % of Diptera with low biological monitoring working party also indicated water ecology deteriorated mainly at midstream sites. Multivariate analysis of classical analysis, canonical correspondence analysis and none metric multidimensional scaling also indicated ecological deterioration at midstream sites mainly at S5, S6, S7 and S8.

**Conclusion:** Midstream was relatively more polluted than upstream due to influence of pollutant from Finoteselam town. The ecological status of upstream segments of the river is very good with slight organic pollutions, and at midstream is poor and significant degree of organic pollutions; whereas the status of the downstream is fair with fairly substantial water pollution showing moderate ecological disturbance. In general, midstream the sampling stations show the deterioration in water ecology and thus necessitated a need for mitigation measure to save the Lahi.

## Introduction

Ecological disturbance is the disturbance of chemical, physical and biological integrity that influence the interdependence of living organism in the environment. When an ecosystem gets polluted, the natural balance in the system is disturbed and this affects the organisms in different ways (Karr, 1991). Rivers are sources of substantial biodiversity and support numerous species from all of the major groups of organisms ranging from microbes to higher forms (MEA, 2005).

Over the past few decades, aquatic ecology has been subjected to a great number of anthropogenic impacts. So that degradation of water resources with pollutant effluent occurs by altering attributes that influence and determines the integrity of surface water resources, such as water quality, habitat structure, flow regime, energy source and biotic interactions that influence the ecological integrity of the system. They are seriously threatening the ecological integrity of most aquatic ecosystems on earth and within this terrible and real scenario, water pollution and its ensuing habitat degradation is the most global challenge in river ecosystems (Camargo, 2017).

As dynamic systems rivers and cities have been in interaction under changing relations over time, and the morphology of many cities has risen through a long and steady struggle between the city functions and the river system flowing inside. This makes river cities an interesting case to study how the presence of geographical features interacts with spatial morphology in the formation of cities (Abshirini and Koch, 2016).

Clean water is a fundamental resource for socio-economic development & transformation; it is essential for maintaining healthy environment and smooth function of ecosytems. There is a raising demand for fresh water resources as a result of increasing population. It has become difficult to treat the current context of growing pollution world-wide. Such problem requires urgent attention, since water is scarce and such an important resource needs detailed scientific research all over the world in order to sustain and protect the water resource from pollution and for its wise utilization. Water resources provide valuable food through aquatic life cycle and irrigation for agriculture production. However, liquid and solid wastes produced by human settlements and industrial effluent disturb most of the water ecology throughout the world.

According to (Giorgio et al., 2016) most evident effects of human pressure on rivers are pollutants like organic residues and heavy metals, acidification and alterations of hydrology and morphology, modification of chemical parameters and variation in biological communities. Activities such as industrialization, urban effluents, deforestation, drainage of wetlands and diffuse sources linked to agriculture have been causing great impact on aquatic ecology in (Prabhakaran et al., 2017, Giorgio et al., 2016 and Al-shami et al., 2011)

According to (Munyika et al., 2014 and Nelson et al., 2016) now a day freshwater ecosystems disturbance is the most threatening environmental effect on earth and its normal functionality, sustainability and their services is declining. Environmental pollution has become a key focus of concern all over the world and has many forms. The air we breathe, the water we drink, and the ground where we cultivate our food crops to health problems and lower quality of life. Among all the environmental pollutants, pollution of freshwater resources especially flowing waters is a matter of great concern. In Africa rapid growth of human populations and the attendant increase in domestic sewage, agricultural development and industrialization are the main causes of water quality as well as ecological deterioration.

According to (Kasangaki et al., 2008) about 30% of the world’s tropical forest is found in Africa, which serves as one of the main carbon sinks. However, since almost all the people in this region (90 %) rely for their energy source on wood, forests are being cleared from time to time this vegetation cover is endangered. On the other hand, land clearing by farmers for agricultural activities may contribute as much as fuel wood gathering in the depletion of tree stocks. As a result; erosion and loss of biodiversity are becoming eminent. In addition, surface water resources are threatened by loss of shade and protection of river banks due to anthropogenic activities.

In Ethiopia, there are many studies done to assess effects of urban pollution on rivers consequently, discharge of untreated effluent from industries, solid wastes and wastewater from households and institutions, are the major sources of pollution of the rivers flowing through the city (Beyene et al., 2009a; 2009b, Dejene Hailu, 1997).

Currently, only 11.4% of Addis Ababa’s population in the urban slums and 41.2% of the city’s total population has access to improved sanitation. Most people in the urban slums (80.4%) used unimproved sanitation facilities and 8.2% practiced open defecation (Abebe Beyene *et al*., 2015).

Before 30 years, (Berhe and Hynes 1989) have found that the macroinvertebrate assemblages in terms of percentage of the Ephemeroptera, Plecoptera and Trichoptera (% EPT) and species richness declined in the sites of the Kebena River located in between Addis Ababa city. Another researcher (Hughes et al., 2009) different biological indicator organisms, which encompasses aquatic organisms like macroinvertebrates, fish, algae, diatoms, macrophytes, riparian and aquatic vegetation and plankton are commonly used as indicators organisms to evaluate the health or pollution status of river ecology.

River water bodies are widely used as disposal sites of solid and liquid waste in the world (Cheng et al., 2018). Discharges of pollutants to the fresh water ecosystem results in a consequent reduction of aquatic life (Souza-Bastos et al., 2017). Thus, the uncontrolled release of waste from urban, domestic, agricultural, and industrial facilities can affect the water quality of the receiving water body (Alemayehu et al., 2005). And this may leads to the severity of ecological disturbance across the river.

According to (Gebre and Van Rooijen, 2009) the main sources of pollution that enters urban surface water bodies such as streams, rivers and reservoirs are discharge of untreated effluents, poor provision of sanitation facilities, rapid population growth, uncontrolled urbanization and improper waste disposal are the major sources of pollution of the rivers flowing through the town.

African countries are facing different challenges on other way towards development. The rapid growth of human populations and the attendant increase in domestic sewage, agricultural development and industrialization are the main causes of water ecology deterioration (Leveque, 1997).

Water quality disturbance from human activities are likely continuing harming human and ecosystem health (Delpla et al., 2009). Available information (MEA, 2005) verifies that excessive nutrient loads and organic pollutant are among the fresh water contaminants of primary concern.

Water pollution profiles of rivers and streams has been undertaken on different rivers and streams in Ethiopia such as; Kebena stream (Berhe, 1988), Great and Little Akaki rivers (Alemayehu et al., 2005), Modjo river (Leta et al., 2003), Sebeta river (Mamo, 2004), Awash river (Bekele,1999) and disturbance of the ecological quality of Awetu River as a result of discharge of municipal wastes and urban runoff has been indicated (Hailu, 1997).

Poor farming methods, destruction of forests, vegetations clearance, discharge of poor or untreated wastes, sand and stone derange, open bathing, car and cloth washing, and poor solid waste management practices which are the most common causes of catchment degradations in the town.

A supply of clean water is an essential requirement for the establishment and maintenance of ecological integrity. Water resources provide valuable food through aquatic life cycle, irrigation for agriculture productions and animal watering. However, improper liquid waste discharge and solid wastes produced by human settlements and industrial activities leads to negative effect to human health and environment as pollution of soil and river water source.

## Objectives

### General objective

▪ To assess the influence of Finoteselam town on ecological status of Lahi river

### Specific objectives

▪ To investigate the physicochemical status of Lahi in the upstream, midstream and downstream from finoteselam.
▪ To examine the abundance of macroinvertebrate assemblage along Lahi river in the upstream, midstream and downstream from finoteselam town.
▪ To identify major anthropogenic activities causing ecological disturbance of Lahi river while crossing finoteselam town.
▪ To assess the physical habitats status at the sampling sites along the river.

### Research questions

▪ Is there a difference among in the upstream, in the middle and downstream of macroinvertebrate assemblage and physicochemical status along Lahi River flowing through finoteselam town?
▪ What are the major anthropogenic activities causing ecological disturbance of Lahi?

## Methods and Materials

### Description of the study area

The study area is located 345 km away from Addis Ababa, the federal capital city of Ethiopia. Finoteselam town is the administrative city for west Gojjam. Lahi river is one of the largest river in finoteselam town

### Selection and description of sampling sites

A total of 11 sampling site coded from S1 to S11 (fig. 1) were chosen along Lahi River and the sampling criteria were leveled of human impact on the river, the study considering processes affecting ecological quality and their influences, accessibility and evaluate the environmental impact of wastes on Lahi River.

**Figure 1.**
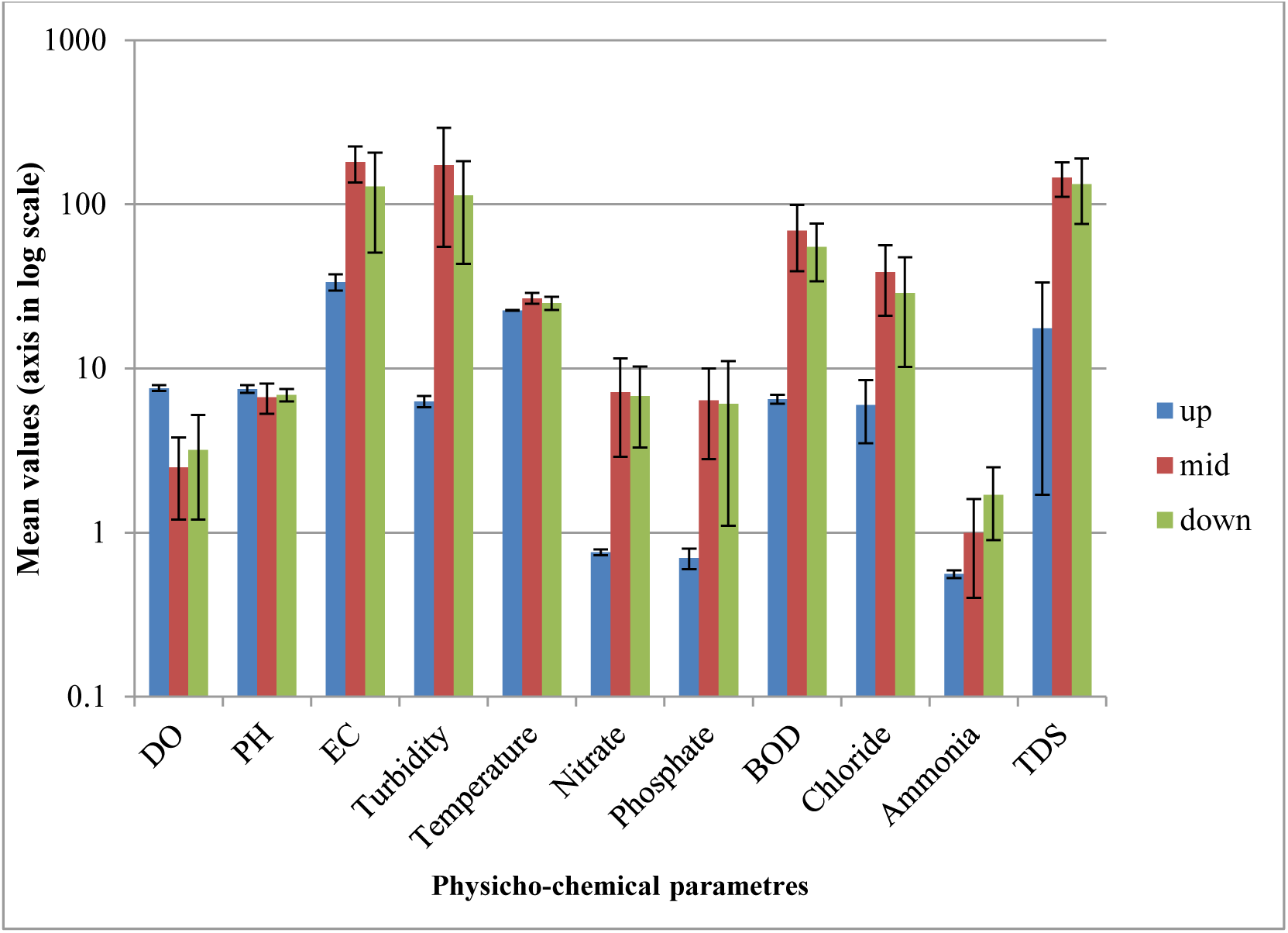
Mean concentraion of physicochemical parameters at threee different stream (Up-Upstream, Mid-Mid-stream and Down-Downstream) along Lahi River.

Sites (S1, S2, and S3) were selected as reference sites before entering the town; using as benchmark to compare changes in other sites. These sites (S1, S2, and S3) were located in the upper part of the river. The other five sites (S4 to S10) were selected on the basis of the prominent land use in the river where as S11 is far from the town. Site (S4, S5, S6, S7 and S8) were midstream and there were extensive human settlement, waste effluent discharge, car washing, and commercial activity, open bathing, cloth washing, sand and stone derange, solid and liquid waste from the town. Here, the water and environment had obnoxious odor when walk across the road. Site (S9, S10 and S11) were downstream, S9 and S10 located in the town, however less human settlement than midstream. At S11 no urbanization seen, this site is far from the town and relatively, low anthropogenic activity to compare the other two downstream sites.

### Study design and period

Cross-sectional study design to assess ecological status of the river from February-1 to April-1 2019

### In-situ measurements

#### Water chemistry

DO, Water pH, conductivity, turbidity, temperature and river discharge were measured at each sampling site with multi meter hand-held probe and turbidity meter (Wag-WT3020) during the sampling periods (Table 2). Altitude, longitude and latitude were measured using global positioning system (GPS).

**Table 1.** Methods used for physicochemical data analysis.

| **Water Physicochemical Parameters** | **Units** | **Methods of analysis** |
| --- | --- | --- |
| **Ph** | Log units | pH meter |
| **Water Temperature** | °C | Hanna Conductivity meter |
| **Electrical Conductivity** | $\mu\text{scm}^{-1}$ | Hanna Conductivity meter |
| **Dissolved Oxygen** | $\text{mgL}^{-1}$ | Winkler method |
| **Turbidity** | NTU | turbidity meter (Wag-WT3020) |
| **Phosphate** | $\text{mgL}^{-1}$ | Stannous Chloride method<br>(UV-Vis spectrophotometer at 690nm) |
| **Chloride** | $\text{mgL}^{-1}$ | Argentometric method (titrating by<br>Ag No3 using K2 Cr O4 as indicator) |
| **Nitrate** | mgL<sup>-1</sup> | Phenoldisulphonic Acid Method<br>(UV-Vis spectrophotometer at 410nm) |
| **Ammonia** | mgL<sup>-1</sup> | Nesslerization method<br>(UV-Vis spectrophotometer at 425nm) |
| **Total Dissolved Solids (TDS)** | mgL<sup>-1</sup> | TDS dried at 180 °C (drying at oven) |
| **Biological Oxygen Demand** | mgL<sup>-1</sup> | Volumetric method |

**Table 2.** Physicochemical and habitat status results along Lahi River.

| Physicochemical parameters and habitat status | Upstream (n=3) |  |  | Midstream (n=5) |  |  |  |  | Downstream (n=3) |  |  |
| --- | --- | --- | --- | --- | --- | --- | --- | --- | --- | --- | --- |
|  | S1 | S2 | S3 | S4 | S5 | S6 | S7 | S8 | S9 | S10 | S11 |
| DO($\text{mgL}^{-1}$) | 7.92 | 7.8 | 7.21 | 4.47 | 2.57 | 3.13 | 1.28 | 1.11 | 3.39 | 1.12 | 5.11 |
| PH | 7.21 | 7.27 | 8.1 | 6.4 | 6.28 | 6.48 | 9.2 | 5.4 | 7.5 | 6.2 | 7.2 |
| EC($\mu\text{scm}^{-1}$) | 30.8 | 32 | 38 | 72.4 | 165.5 | 160.3 | 167.7 | 188.2 | 145.3 | 197.1 | 43.8 |
| Turbidity (NTU) | 5.7 | 6.8 | 6.3 | 27 | 195.3 | 192.6 | 174.3 | 280.6 | 135 | 160 | 45.2 |
| Water T($^{\circ}\text{C}$) | 22.4 | 22.6 | 22.8 | 24 | 27 | 26 | 28 | 29.3 | 24.5 | 28.1 | 23.7 |
| Velocity (m/s) | 1.25 | 1.3 | 1.1 | 0.9 | 0.71 | 0.81 | 0.53 | 0.5 | 0.61 | 0.58 | 0.75 |
| Rivedischarge($\text{m}^3/\text{s}$) | 11 | 10.6 | 8 | 5.9 | 1.7 | 2.67 | 0.9 | 0.7 | 2.2 | 1.8 | 6.5 |
| Nitrate ($\text{mgL}^{-1}$) | 0.74 | 0.75 | 0.8 | 2.94 | 4.55 | 5.39 | 9.64 | 13.56 | 8.12 | 10.7 | 1.63 |
| Phosphate ($\text{mgL}^{-1}$) | 0.64 | 0.73 | 0.88 | 1.71 | 6.75 | 4.14 | 8.5 | 11.1 | 7.57 | 9.7 | 0.97 |
| BOD ($\text{mgL}^{-1}$) | 6.1 | 6.6 | 6.95 | 31.4 | 66.7 | 52.6 | 86.2 | 109.35 | 61.65 | 67.4 | 36.7 |
| Chloride ($\text{mgL}^{-1}$) | 3.3 | 6.6 | 8.3 | 9.9 | 46 | 41 | 46 | 50.8 | 38 | 43.3 | 7.8 |
| Ammonia ($\text{mgL}^{-1}$) | 0.56 | 0.54 | 0.6 | 0.7 | 0.7 | 0.66 | 0.9 | 2.29 | 2.11 | 2.4 | 0.83 |
| TDS ($\text{mgL}^{-1}$) | 8 | 8.8 | 36 | 88 | 156 | 142 | 164 | 178 | 136 | 189 | 74 |
| Canopy cover(%) | 90 | 70 | 65 | 10 | 0 | 0 | 0 | 0 | 0 | 0 | 50 |
| Habitat score | 173 | 151 | 116 | 136 | 47 | 73 | 61 | 43 | 73 | 44 | 116 |

Note: habitat score, as optimal (160-200), suboptimal (100-159), marginal (60-99) and poor (<60).
Kruskal-Wallis test indicated that the mean values of dissolved oxygen , electrical conductivity, temperature, turbidity, phosphate , nitrate , chloride , BOD,TDS, river discharge ,canopy cover and habitat score significantly differed among upstream, midstream and downstream ( $p<0.05$ ). Other water quality parameters such as pH, ammonia and surface velocity were not significantly ( $p>0.05$ ) differed among upstream, midstream and downstream water level.

#### Physical habitat

Habitat conditions information at each site were collected via visual estimation with measured of the physicochemical parameters and recorded for each monitoring reach during collection of macroinvertebrates sample. Habitat features were scored with the UPA”s rapid bio assessment protocol the procedure given in Barbour et al. (1999). Habitat assessment involved rating 10 habitat parameters (epifaunal substrate/available cover, embeddedness, velocity/depth regime, sediment deposition, channel flow status, channel alteration, frequency of riffles, bank stability, vegetative protection and riparian vegetative having habitat score classifying each sampling site as optimal (160-200), suboptimal (100-159), marginal (60-99) and poor(<60). All the parameters were evaluated and rating on a numerical scale from 0-20 for each sampling reach at each site. Depth and river velocity were measured using dip stick and flotation methods, respectively. Then the rating then total and expressed as score to provide the final habitat quality ranking (Table 3).

**Table 3.** Relative abundance of macroinvertabrate assemblages for the three stream category

| Family | Stream Category |  |  |
| --- | --- | --- | --- |
|  | Upstream | Midstream | Downstream |
| Gompidae | 39 | 7 | 0 |
| Aeshnidae | 48 | 6 | 0 |
| Coenagrionidae | 7 | 4 | 2 |
| Velidae | 12 | 4 | 5 |
| Perlidae | 2 | 0 | 0 |
| Batidae | 4 | 2 | 3 |
| Gyrinidae | 8 | 0 | 2 |
| Notoectidae | 7 | 0 | 5 |
| Corixidae | 3 | 2 | 8 |
| Chironomidae | 6 | 35 | 18 |
| Belostomatidae | 8 | 8 | 4 |
| Heptagenidae | 8 | 0 | 0 |
| Caenidae | 14 | 0 | 0 |
| Hydrophilidae | 6 | 0 | 6 |
| Hydropsychidae | 11 | 1 | 0 |
| Physidae | 0 | 136 | 37 |
| Libellulidae | 11 | 0 | 0 |
| Elmidae | 7 | 2 | 0 |
| Nacuridae | 8 | 8 | 0 |
| Chloroperlidae | 9 | 0 | 0 |
| Calopterygidae | 13 | 0 | 0 |
| Planorbida | 0 | 30 | 30 |
| Lymnaeidae | 0 | 17 | 16 |
| Ceratopogonidae | 13 | 0 | 0 |
| Tipulidae | 0 | 0 | 3 |
| Oligochaeta | 0 | 0 | 6 |
| Culicidae | 0 | 6 | 4 |
| Tabanidae | 0 | 0 | 3 |
| Psychomyiidae | 14 | 0 | 0 |
| Athericidae | 11 | 0 | 0 |

#### Anthropogenic activity assessment

The assessment of human activity were used by observed and recorded each activity including the portion of the sampling site 100m surrounding station (Barbour et al. 1999) and considering based on the major human activities in the area.

### Sampling collection and protocol

#### Water samples

Composite water sample were collected by 1-L polyethylene sampling bottles and10 cm below the surface as indicated in APHA et al. (1999). At each sampling site, the bottles were rinsed at least three times before sampling.

#### Macroinvertebrate samples

Macroinvertebrate samples from the sampling sites concurrently were collected with the water sampling during stable flow. According to international standardized protocol using a triangular D-frame kick hand net with a 250 μm mesh size in all the available habitat types (multi habitat sampling procedure), such as riffles, macrophytes, pools and bedrock collectively for 5 min per site (Barbour et al., 1999). At each sampling site, macroinvertebrates were taken by submerging the hand net in the river at different depths, sweeping and kicking at each location, covered both vegetation and open-water areas to incorporated habitat heterogeneity and benthic substratum were dislodge to any attached macro invertebrates (Berger et al., 2017). After the macroinvertebrates sampling were collected and transferred into a bowel and washed with sufficient amount of water were added and the supernatant was poured onto a sieve to retain the macroinvertebrates while removed the mud. This was repeated until all the macroinvertebrates separated from mud. Then preserving in 95% ethanol

After all samples collected then placed in an insulating ice packs cooled boxes and transported to BDU Environmental Sciences and Technology laboratories with due care for chemical and biological analyses and water samples stored in a refrigerator at 4°C until analysis

### Laboratory analyses

#### Nutrients loads

From each water sample nitrate, phosphate, chloride, TDS, ammonia and BOD were analyzed by following the procedures outlined in APHA et al. (1999) to determined level of ecological disturbance of nutrient. The methods were presented in Table 2.

#### Macroinvertebrate identification

For all macroinvertebrate samples were transferred in to Petri dish in order to easily observed and peaked up the organisms with forceps then sorted and identified the specimen at family level used a stereo dissecting microscope and samples of macroinvertebrate were carried out based on aquatic taxonomic identification keys level, given in (Barbour et al. 1999) then based on by their specific category of all specimens counting and putting in one vial.

### Data analysis

Significances of ecological deterioration variability of the water quality parameters and macroinvertebrate assemblage were tasted by using Kruskal-Wallis multiple comparison test and Pearson correlations were performed between bio-assessment indices and physic-chemical variables to determine the sensitivity of each index to specific variables. Statistical analyses were performed using Excel and Statistical Package for Social Sciences (SPSS) version 24. Cluster Analysis, and Canonical Correspondence Analysis were done using Paleontological Statistics software package for education (PAST 3) version 3.18. In addition, to compare the three streams post-hoc comparisons were made with the STATISTICA^®^ software package version 7.1 as well as non-metric multidimensional scaling to discriminate irrelevant macroinvertebrate metrics for detection of human impact was performed using the R statistical package (R Development Core Team, 2008).

#### Shannon Diversity Index

Shannon diversity index (Shannon, 1948) is the most used metric which measure different aspects of diversity for macroinvertebrates to describing the species abundance of individuals within the eleven sites were calculated based on (Shannon, 1948). The index was calculated from the proportional abundances of each species as with the formula.

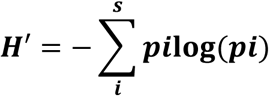

Where,

H’ is the standard symbol for the maximum Shannon index, and pi is the proportion of i’th species.

*i* = an index number for each species present in a sample.

#### Evenness

Evenness was calculated for macroinvertebrates as the ratio of diversity with the maximum possible diversity for the number of species found as:

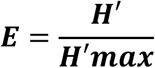

Where, H’ is Shannon index and H ‘max = maximum possible Shannon’s diversity

### Community Loss Index

CLI measures the loss of benthic tax in a study site with respect to a reference site. Value range from 0 to “infinity” and increase as degree of dissimilarity between the sites increase (Mandaville, 2002). The index was calculated base on the following formula.

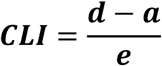

Where:

“a” is the number of taxa common to both sites, “d” is the total number of taxa present in the reference site, and “e” is the total number of taxa present in the study site.

### EPT richness/EPT index

EPT index displays the taxa richness within the insect groups which are considered to be sensitive to pollution, and therefore should increase with increasing ecological water quality. The EPT index is equal to the total number of families represented within these three orders in the sample.

### Family Biotic Index

FBI is an average of tolerance values of all the macroinvertebrates families in a sample (Hilsenhoff, 1988). FBI was calculated by multiplying the number in each family by the tolerance value for that family, summing the products, and dividing by the total macroinvertebrate in the sample. The family-level tolerance values range from 0(very intolerant) to 10(highly tolerant) based on their tolerance to organic pollution. The FBI was then used to evaluate the pollution stats of the water for each sampling sites and the three streams by comparing with the standard used to rate the ecological water quality status. The index is calculated based on the following formula.

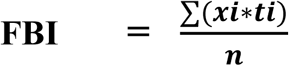

Where ***xi*** is abundance of taxon ***i, ti*** is the tolerance value of taxon ***i*** and ***n*** is abundance in the sample.

### Biological Monitoring Working Party

BMWP score is an index for measuring the biological quality of rivers using species of macroinvertebrates as biological indicators at the family level score which is representative of the family’s tolerance to water pollution (Walley and Hawkes, 1997). Each family is then given a score between 1 and 10. Tolerant (1) and intolerant (10). The overall BMWP Score for a site is the sum of all of the scores of each family present at that site. Then compare with the standard. If the value greater than 100 is associated with clean rivers, whilst heavily polluted rivers score less than 10.

### Cluster Analysis

Cluster analysis one which is an exploratory multivariate data analysis tool that sorts different objects into groups in a way that the degree of association between two objects is maximal if they belong to the same group and minimal otherwise. This helps to arrange the data into a meaningful structure (Quinn and Keough, 2002). Different clustering algorithms like single linkage, complete linkage, Ward’s method, etc. can be applied. In this thesis, hierarchical clustering of macroinvertebrate and physicochemical data were performed using Bray-Curtis distance and Ward’s clustering algorithm to assessing their effectiveness in classifying sample sites based on pollution load and sources of pollution. The Paleontological Statistics software package for education (PAST 3) version 3.18 was used to undertake this cluster analysis.

### Non-metric multidimensional scaling

Non-metric multidimensional scaling was performed to the similarity between samples and macroinvertebrate metrics from the reference condition. This method is vital for the development of bio-monitoring metrics for one region (Paul *et al*., 2005). NMS is not an eigen-value-eigenvector technique like principal component analysis or correspondence analysis. Because of this, an NMS ordination can be rotated, inverted, or centered to any desired configuration. NMS makes no assumption of linear relationship, so it is well suited for a wide variety of data. NMS allows the use of any distance measure of the samples. This makes NMS suitable for analysis of samples and variables and for choosing appropriate variables during the metric development. To discriminate irrelevant macroinvertebrate metrics for detection of human impact was performed using the R statistical package (R Development Core Team, 2008).

### Canonical Correspondence Analysis

The analysis of environmental and biological data, canonical correspondence analysis (CCA) is used to assess the relationship between the two groups of variables. CCA can be applied to detect both species-environment relations, and for investigating the response of species to environmental variables. CCA constructs linear combinations of environmental variables, along which the distributions of the species are maximally separated by eigen values produced which are measured by CCA (Shaw, 2003).). In this case, CCA is one of the most used ordination technique in ecology. The PAST for Windows Version 3.8 software package was used to undertake the CCA analysis.

## Result

### Physicochemical parameters and habitat status

The physicochemical parameters of Lahi River were summarized into mean, standard deviation, minimum and maximum values among upstream, midstream and downstream sites (Fig.5). From the upstream sites, the oxygen level was above 7 mgL^−1^, while turbidity was increasing in the downstream direction where the maximum was at midstream sites. Likewise, BOD was higher in the midstream sites which showed recovery in the downstream sites. The physical habitat evaluation scores at the sampling sites ranged from 43 at S8 to 173 at S1. Relatively the highest score was recorded at upstream sites whereas the lowest score was recorded at midstream sites. Table 3 shows the detail values of physico-chemical parameters and habitat status.

**Figure 2.**
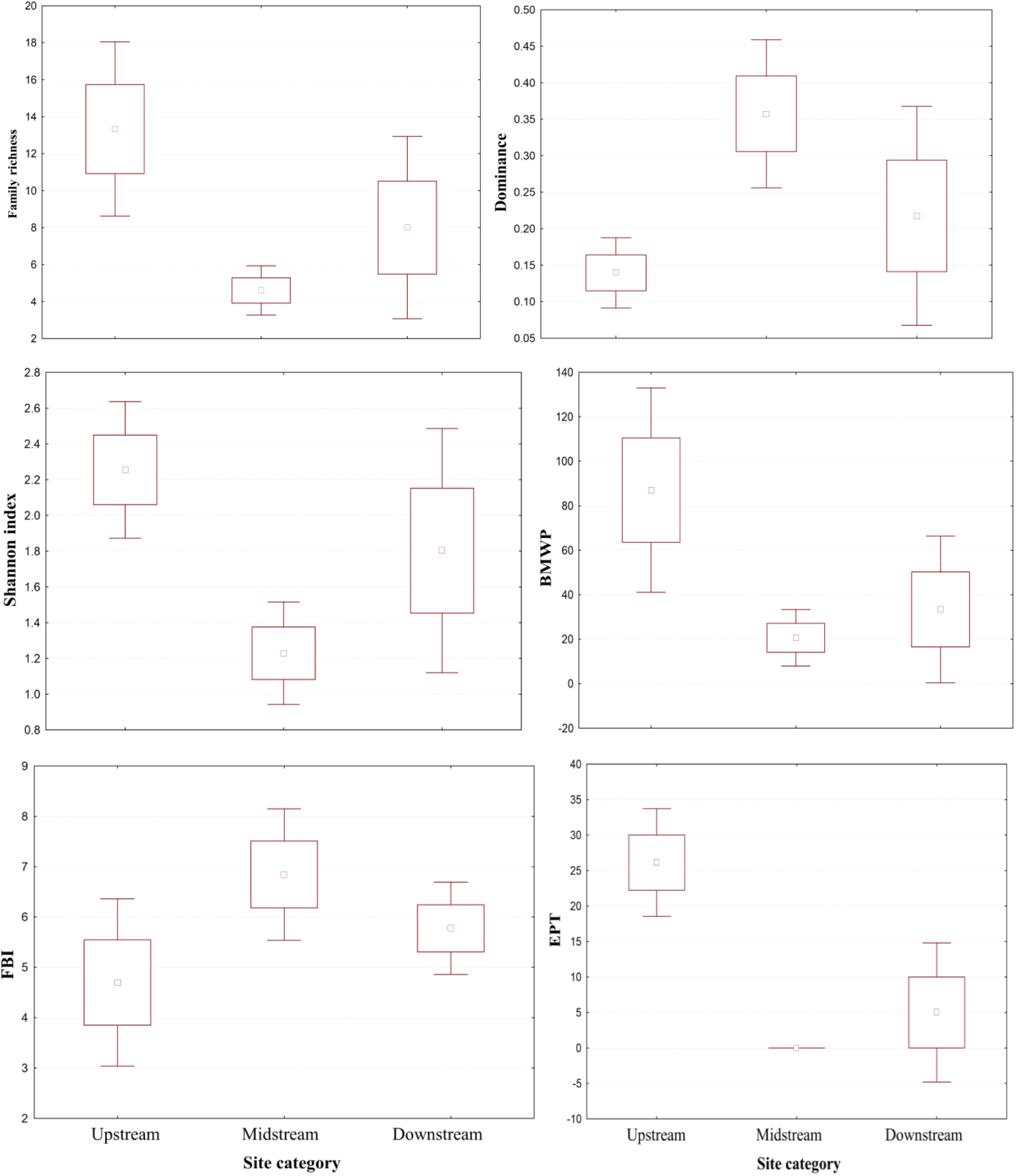
Box (standard err) and whisker (95%) confidence interval) plots of macroinvertebrates in upstream, midstream and downstream sites from the town.

**Figure 3.**
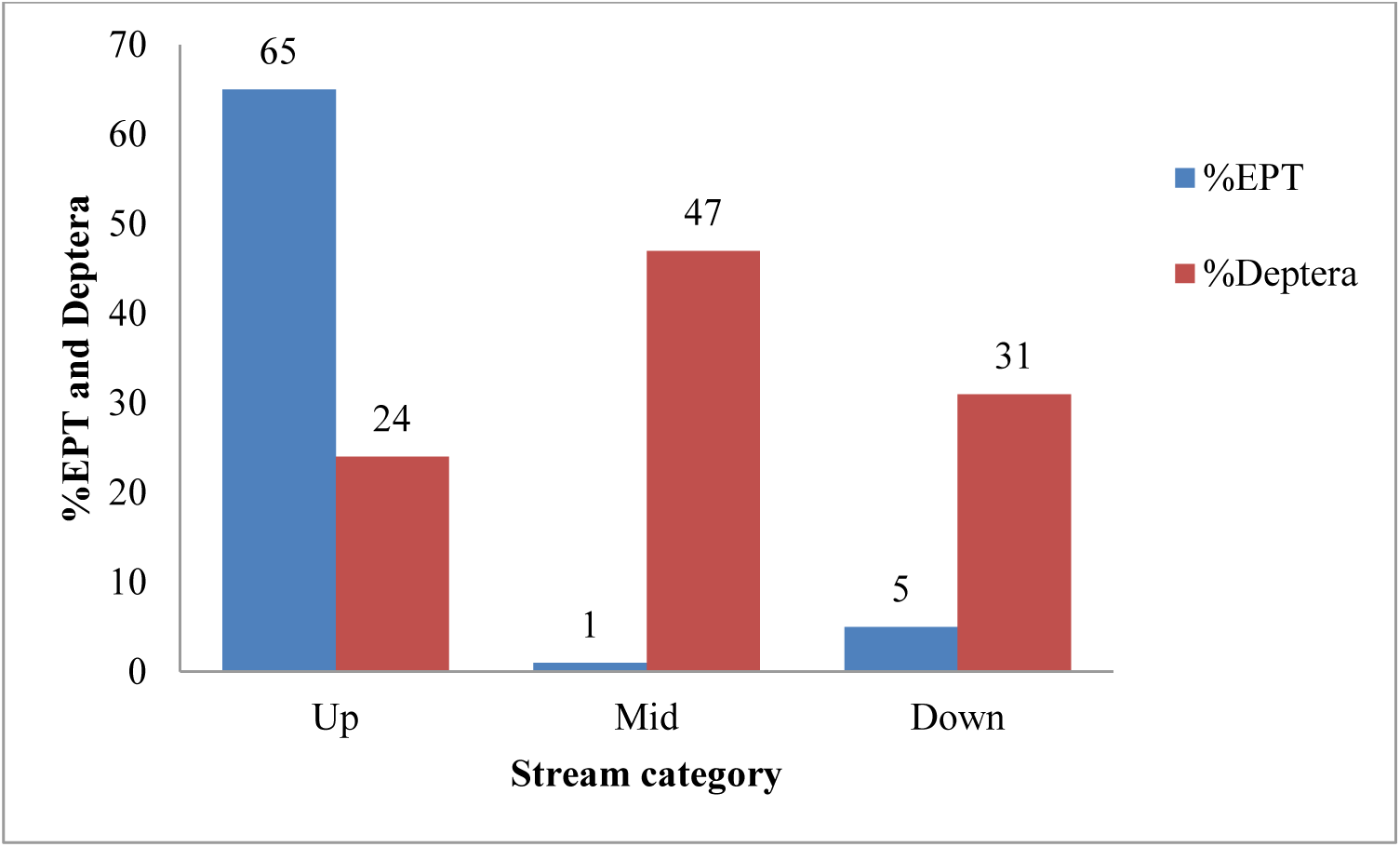
Both %EPT and %Deptera at the three different streams (Up-Upstream, Mid-Midstream, Down-Downstream) along Lahi River

**Figure 4.**
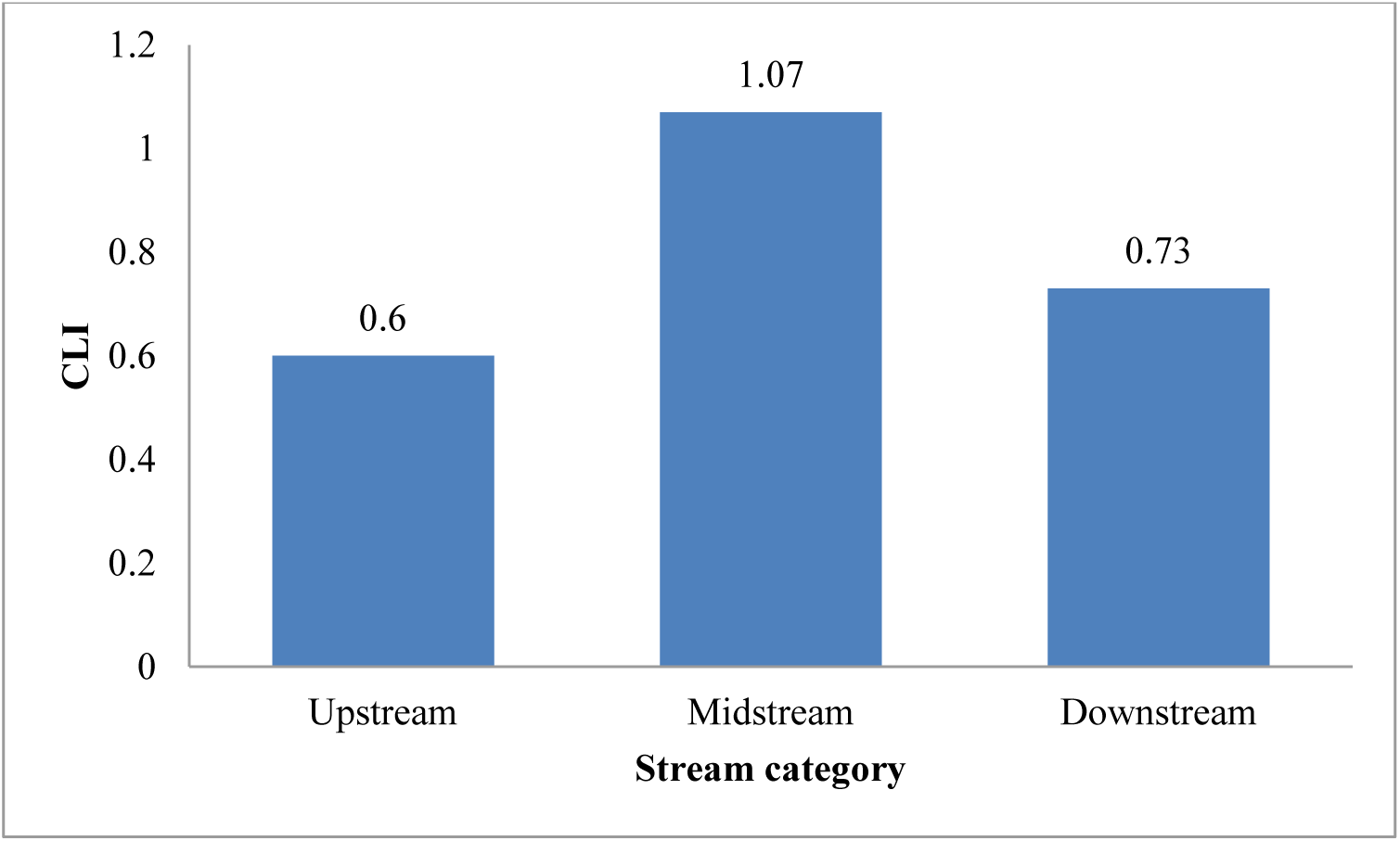
Ecological status variation in CLI at upstream, midstream and downstream along Lahi River

**Figure 5.**
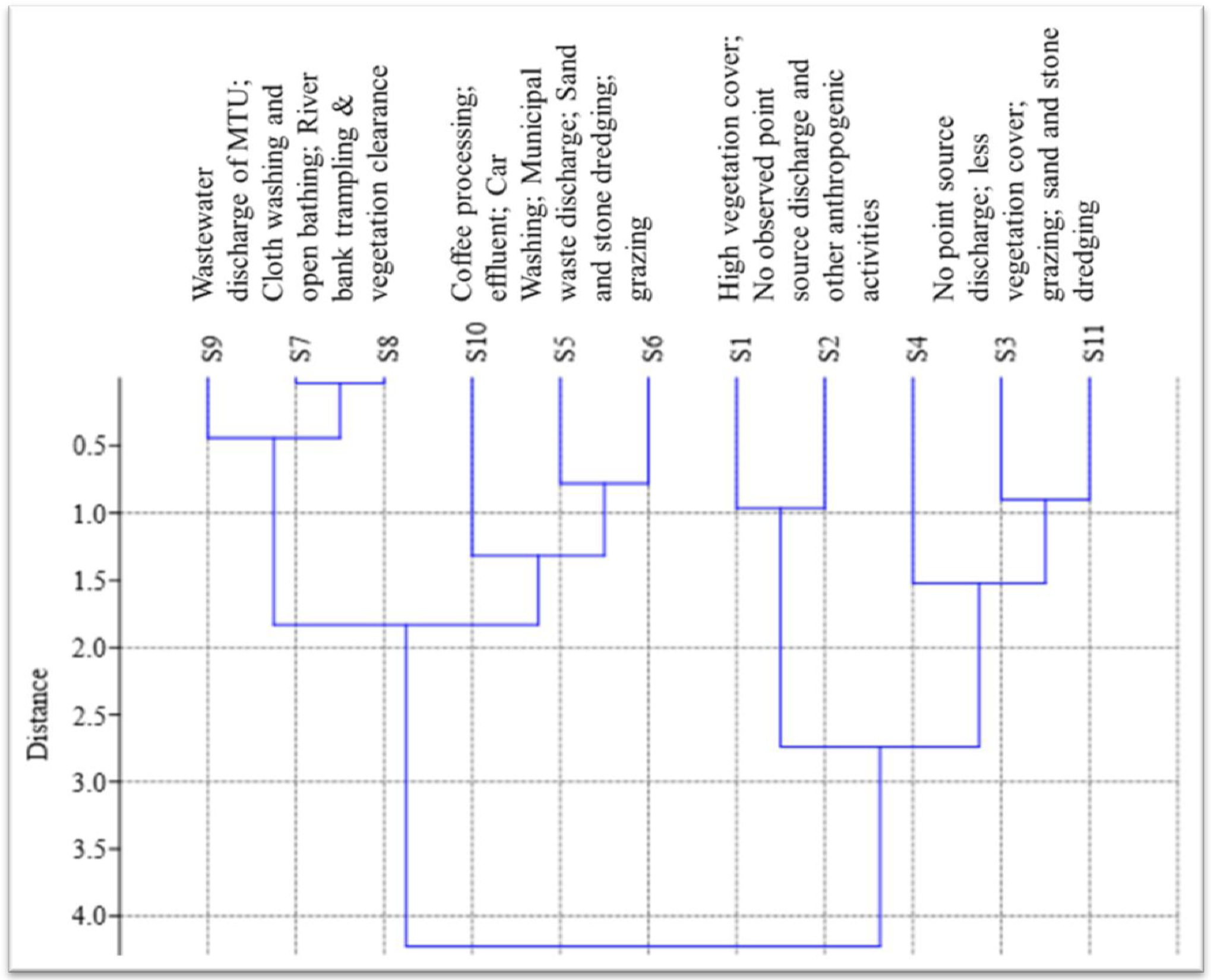
Hierarchical clustering macroinvertabrates data using Ward’s clustering algorithm and human activities within the sites.

#### Macroinvertabrate assemblages

A total of 30 taxa with 746 individuals of benthic macroinvertabrates were collected from all sampling sites in which the highest (19 families comprising 172 individuals) and the lowest (3 families comprising 20 individuals) taxa richness was collected at S1 and S5, respectively. Upstream has significantly better macroinvertabrates assemblage (23 taxa) than midstream (13taxa) (p-value<0.05) and downstream (19 taxa).

*Batidae* was common taxa in the three stream while *Phsidae and Planorbidae* were collected only in the midstream and downstream exclusively S11. However *Perlidae, Caenidae, Athericidae, Psychomyiidae, Chloroperlidae, Libellulidae, Heptagenide* as well as *Calopterygidae* were founded only in the upstream sites. From the eleven sites *Physidae* was the most dominant (136) which were mostly collected at mid-stream where as no one snail collected at upstream sites. Generally, taxa richness was tended to declining in the downstream direction where the minimum was at midstream sites. The result presented in Table 4

**Table 4.** Ecological status in the indices analysis results from the sampling sites along Lahi River.

| Indices | Sampling sites |  |  |  |  |  |  |  |  |  |  |
| --- | --- | --- | --- | --- | --- | --- | --- | --- | --- | --- | --- |
|  | S1 | S2 | S3 | S4 | S5 | S6 | S7 | S8 | S9 | S10 | S11 |
| Taxa richness | 19 | 12 | 10 | 7 | 3 | 5 | 4 | 4 | 5 | 6 | 13 |
| Individuals | 172 | 63 | 41 | 23 | 20 | 24 | 83 | 145 | 41 | 74 | 60 |
| Shannon_H' | 2.62 | 2.16 | 1.71 | 1.97 | 1.04 | 1.39 | 0.89 | 1.09 | 1.27 | 1.66 | 2.46 |
| Dominance_D | 0.09 | 0.15 | 0.17 | 0.2 | 0.36 | 0.29 | 0.52 | 0.39 | 0.35 | 0.2 | 0.09 |
| Evenness-E | 0.77 | 0.72 | 0.72 | 0.79 | 0.95 | 0.8 | 0.61 | 0.74 | 0.71 | 0.88 | 0.9 |
| EPT richness | 51 | 10 | 3 | 1 | 0 | 0 | 0 | 0 | 0 | 0 | 5 |
| FBI | 2.8 | 3.2 | 5.39 | 4.56 | 8.5 | 6.48 | 7.72 | 6.96 | 5.65 | 6.64 | 5.03 |
| BMWP | 89 | 84 | 48 | 45 | 13 | 23 | 11 | 11 | 16 | 17 | 67 |
| %EPT | 33.3 | 25 | 20 | 0 | 0 | 0 | 0 | 0 | 0 | 0 | 15 |
| % Deptera | 11.1 | 16.6 | 22 | 14.2 | 66.6 | 40 | 25 | 30 | 20 | 33.3 | 15.3 |
| CLI | 0.73 | 1.16 | 1.4 | 2 | 4.6 | 2.8 | 3.5 | 3.5 | 2.8 | 2.3 | 1 |

Based on Kruskal-Wallis multiple comparison test, the total taxa richness significantly differed (p-value <0.05) among upstream, midstream and downstream. The metric negatively correlated with Nitrate, Phosphate, BOD, EC and turbidity.

### Indices

The Shannon diversity index varied among the sampling sites. The lowest value (0.8) was at S7 while the highest value (2.6) determined at S1, and Evenness exceeded at S5 (Table 5). When species composition was compared, diversity was highest at upstream with slight difference from downstream and lowest at midstream. Although midstream sites are least diverse, its evenness is comparable with upstream. Though in downstream sites are more diverse than midstream and less diverse than upstream sites.

EPT richness was zero at S5, S6.S7, S8, S9 and S10 as no specimen was collected whereas relatively the highest level was recorded at S1 (Table 5). When the three stream are compared with respects to their EPT richness values at midstream is the least value (1%) where upstream (65%) (Fig.7)

Helsenhoff FBI was used to assess the pollution status of the river using macroinvertebrates and the result is presented in Table 5. FBI varied from 2.8 at S1 to 8.5 at S5 among the sites where specimens were collected and relatively the highest value recorded at mid-stream sites (Fig.6). When the three stream categories are compared with respects to their FBI mean values, upstream value (3.7), followed by downstream (5.7) and midstream (6.8).

**Figure 6.**
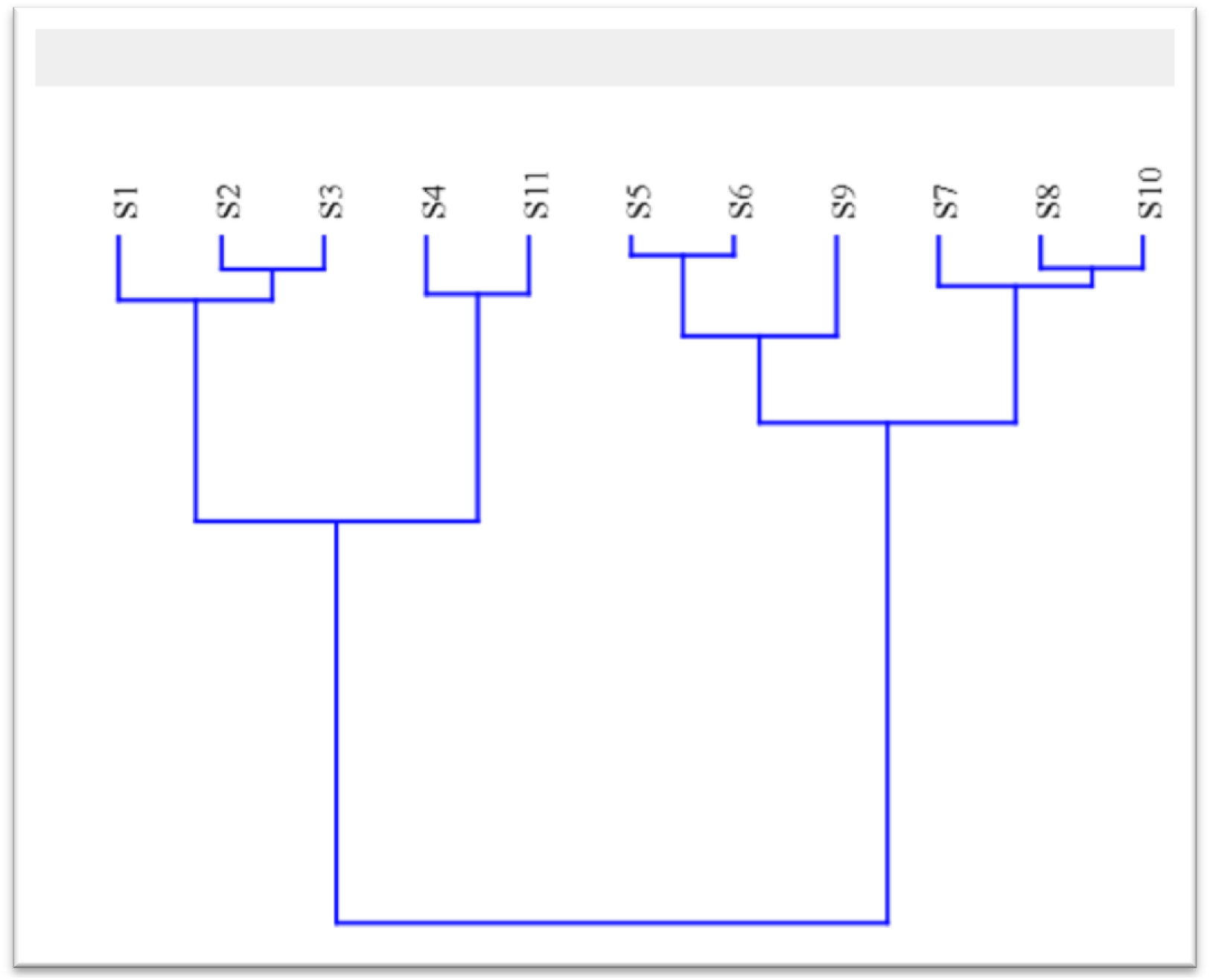
Hierarchical clustering of physicochemical data using Ward’s clustering algorithm

**Figure 7.**
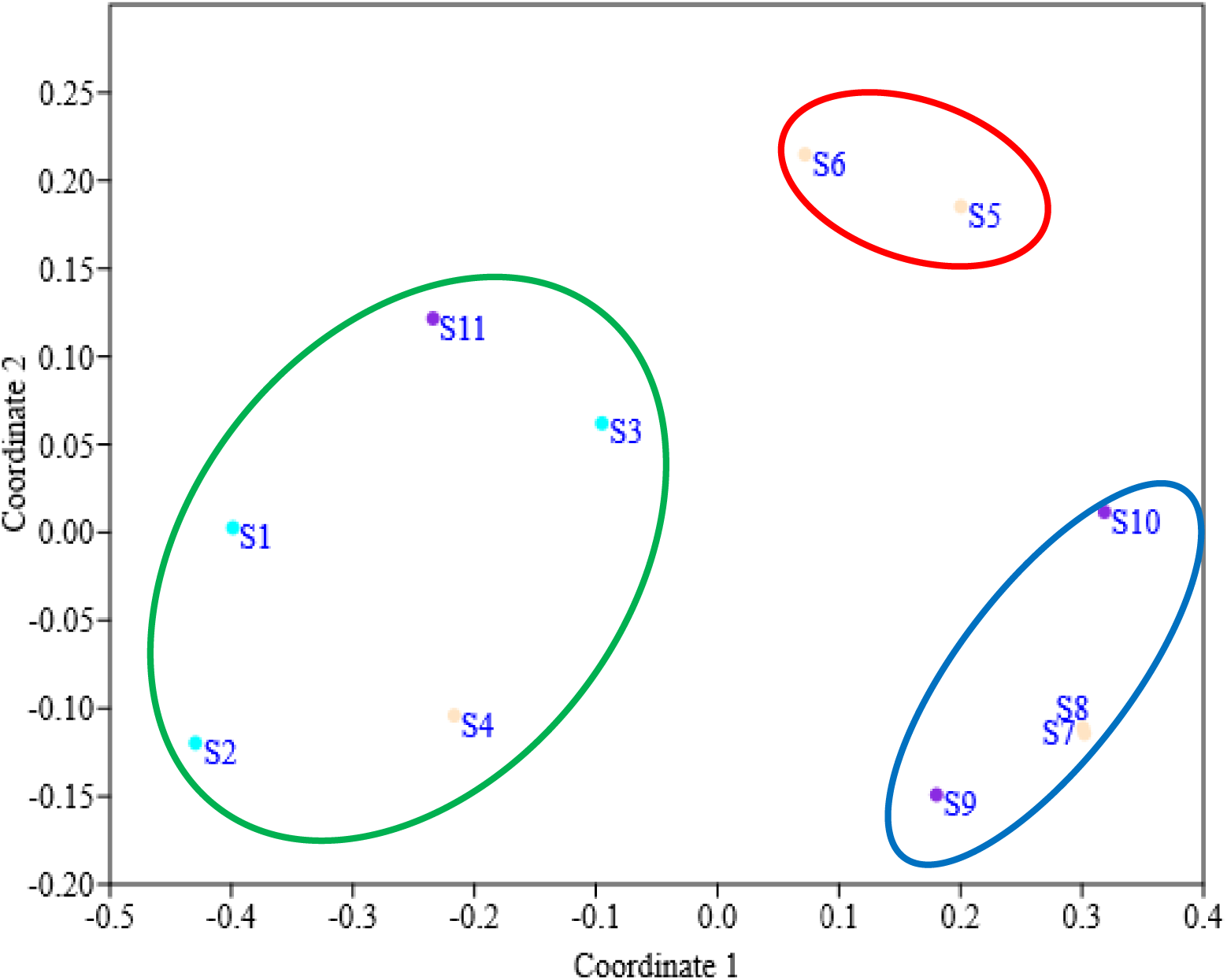
None metric multidimensional scaling plots using macroinvertabrates along Lahi River.

The percentage of *Ephemeroptera Plecoptera Trichoptera* varied from 0% at S5, S6, S7, S8, S9 and S10 to 36.8% at S1 among the sampling sites where specimens were collected and relatively lower at all sites except at S1. The difference in the %EPT taxa among the stream category was somewhat higher in upstream(65%)than midstream(1%).

BMWP score ranged from 11at S8 to 89 at S1 from the sites (Table 5). Among the three stream categories are compared with respect to their BMWP mean values, midstream is the least with value of 20.6, followed by downstream with 43.3 and upstream with 73.6.

Based on Kruskal-Wallis multiple comparison test, the total family richness, BMWP and %EPT indices significantly differed (p-value <0.05) from the midstream sites but not from the downstream sites.

*Deptera* was collected at S5, S6, S8 and S10 whereas no specimen at S1 and S2. *Deptera* comprising mainly *Chironomidae* and *Culicidae* was abundant group at midstream value (47%). Lower %EPT and higher %Deptera were collected at midstream sites as shown the following Figure 7.

Macroinvertebrate CLI was relatively higher at S8 and lower as S1 (Table 5). CLI values, midstream is the highest with value of 1.07; followed by downstream with 0.7 and up-stream with 0.6 (Fig.8). CLI indices upstream significantly differed (p-value <0.05) from the midstream sites but not from the downstream sites (p>0.05)

**Figure 8.**
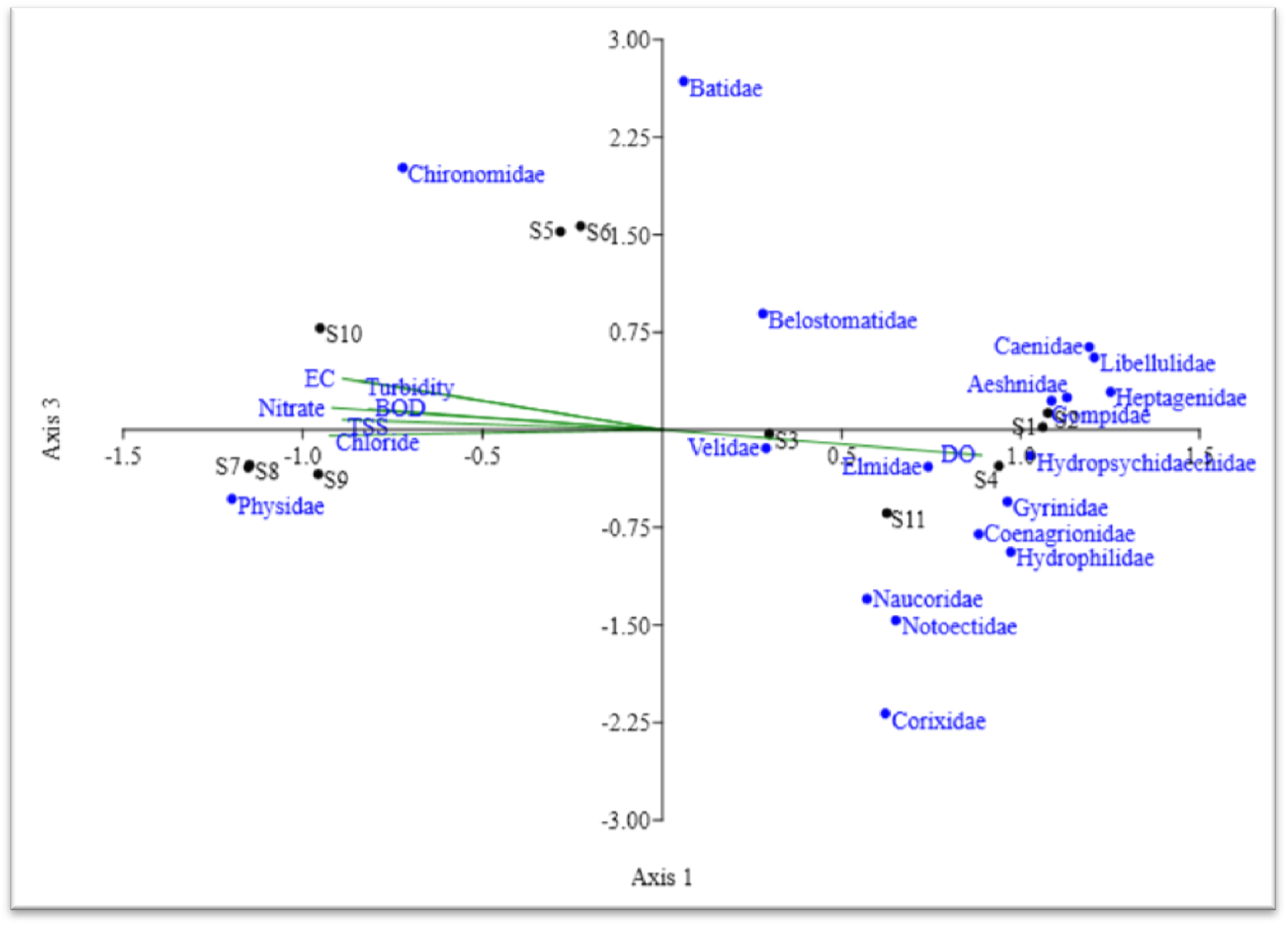
1^st^ and 3^rd^ CCA axes of benthic macroinvertabrates taxa environmental variables and the corresponding sampling sites at Lahi River.

### Multivariate Analysis

Cluster analysis using benthic macroinvertabrate data groped S1 and S2, S3 and S11, S5 and S6 where sampling resulted in low taxa collected (i.e. S7 and S8) in separate clusters but placed S4, S9 and S10 in their own groped (Fig.9)

Cluster analyses using physicochemical data groped S2 and S3, S5 and S6, S8 and S10 as well as S4 and S11 in four different clusters while S1, S7 and S9 were placed separately in their own groups (Fig. 10)

None metric multidimensional scaling plots using macroinvertebrate data. Three major groupings, where sites circled in green (S1, S2, S3, S4 and S11) are with better macroinvertabrates assemblage, in red (S5 and S6) are impacted sites, and while in blue circle (S7, S8 and S10) are those receiving point source discharges (Fig. 11)

CCA triplot for benthic macroinvertabrates indicated similarity between S1 and S2, S7 and S8, as well as among S3, S4 and S11 where as strong similarity between S5 and S6 in ecological water quality. The triplote further indicated a relatively the difference in water quality at S10.

Axis 1 CCA triplote macroinvertabrate data clearly separated S1, S2, S3 and S11 from the other sampling points while Axis 3 separated S5and S6, S7and S8 from other indicating that the sampling sites were different from other sampling sites in terms of water quality. The direction proportional influence of BOD, EC, TDS, Nitrate, Chloride and Turbidity pointing S5, S6, S7, S8, S9 and S10. The directions proportional influence of DO were pointing S1, S2 S3 S4 and S11.

The arrow of environmental variables points in the direction of maximum change in the values of associated variable, and the arrow length is proportional to this maximum rate of change. Physicochemical parameters in the CCA tri plots pointing away from S1, S2, S3, S4 and S11 since the measured values of those variables that increase with increase in pollution load were relatively lower at these sites.

## Discussions

Clean water is an essential requirement for the establishment and maintenance of ecological integrity (Berger et al., 2017). Water resources provide valuable food through aquatic life cycle, irrigation for agriculture productions and animal watering (Bostanmaneshrad et al., 2018). However, improper liquid waste discharge and solid wastes produced by human settlements and industrial activities leads to negative effect to human health and environment as pollution to river water source (Ambelu et al, 2013).

BOD, an indicator of pollution, was greater in all midstream sites, and the highest concentration (109.3 mgL^−1^) was recorded at S8. This high BOD may have been due to the sites taken sewage discharge, and disposal of solid and liquid wastes from finoteselam town. It is quite interesting to note that the DO is less than 3 mgL^−1^ in all the midstream sites except S4. Therfore, statically significant difference was observed for among streams (p<0.005). This may be attributed to poor treated waste discharged from animal waste and urban sewage discharge in the area.

As implicated by U.S. Environmental Agency, DO>5mgL^−1^ is considered favorable for growth and activity of most aquatic organisms; DO<3mgL^−1^ is stressful to most aquatic organisms, while the standard for surface water of Ethiopia ≥4mgL^−1^. Thus all the sampling station in midstream showed higher BOD values compared to upstream and downstream. This reveals clearly that midstream was experiencing a higher level of pollution than upstream and downstream (Fig.5). The higher level of BOD and lower DO occur at midstream sites might be due the introduction of solid and liquid wastes from the town. When the mean concentration of DO measured at the three study stream sites is compared according to the standard set above, the concentration at midstream and downstream sites (2.5 and 3.2 mgL^−1^) respectively. This indicates that the water at midstream and downstream sites are unfavorable for aquatic life, while the mean concentration of at upstream sites (7.6 mgL^−1^) are considered to be favorable for aquatic life.

The concentration of nitrate was grater the critical values in the entire sampling site except upstream sites and from downstream S11. Nitrate concentration of unpolluted surface water seldom exceeds 0.1 mgL^−1^ and whenever it has above 0.2mgL^−1^ enhances eutrophication-ficaion (Chapman, 1996). However, The required amount of nitrate in water for animal drinking use is set as 100mgL^−1^ and, for irrigation water, FAO recommended maximum concentration of 30mgL^−1^ and standard a concentration below 5mgL^−1^ poses either in plant or soil (Wang et al, 2017), whereas the ambient standard for Ethiopia surface water Quality Guide lines (< 50 mgL^−1^). The intense damping of various solid and liquid wastes from the town as well as the effluent discharge point from town in midstream sites (Table 3) could have been the major source of nutrients. The downstream sites, S10 has also considerable nitrate concentration (10.7mg^−1^L) which is perhaps due to the highest nitrate load from midstream and the site receive sewage discharge from the surrounding community. The major anthropogenic sources of nitrate in aquatic ecosystem are sewage, fertilizers, and waste from domesticated animals (Ambelu et al., 2013). The concentrations of nitrate further down at S11 (1.6) the site might be partly attributed to the nutrient filtering effect of riparian vegetation grass and river self purification further down the nitrate loads. Similarly, the concentration of phosphate was much higher than the critical value. According to (Awoke et al, 2016) maximum allowable concentration of phosphate in irrigation water is 2 mgL^−1^ in all most all site except upstream and S11and this may be due finoteselam town that domestic waste discharges, and phosphates detergent to the sites through point and non point source in addition to fertilizers run from the catchment. It has been reported that rapid urbanization, waste discharge, and other anthropogenic activities are known causes of nutrient enrichment and threat river deterioration (Luo et al., 2017)..

PH is an important variable in ecological water quality assessment as it influences many biological and chemical processes within the water body (Affairs, 1996) the pH of most raw water source lies within the range of 6. 5-8.5 and influenced by various factors and processes including temperature, discharge of effluents, runoff, acidic perception, microbial activity and in line with the present finding. The highest pH was recorded at S7 (9.2) might be explained due to waste discharge, cloth washing and open bathing a lot of students statically not significant (P>0.05). Electrical conductivity is measure of the ability of water to electric current. It is related to provide a measure of the total dissolved solids. The rises and/or falls of electric conductivity are attributed to the dissolved solids in water (Colin et al., 2017). Therefore, the TDS contents are directly related to the electrical conductivity (Fierro et al, 2018). This fact holds true for upstream, midstream and downstream sites where the change in TDS is directly related the change in electrical conductivity. The value of TDS much higher than the critical value of animal drinking is required between 100-150 mgL^−1^ (Affairs, 1996)) in all most all sites except at all upstreamsites, S4 and S11 this might be attributed to effluents discharged from various sources in finoteselam town. It has been indicated that conductivity can be measured pollution zone of water, example effluent around discharge (Iwasaki et al, 2018). Turbidity is a measure of cloudiness of water, which is material suspended (floating) in water. Therefore, it is an indicator of the concentration of suspended sediments in water (Riley, 2008). The maximum turbidity recorded at midstream might be attributed to highest sediment lodes through surface runoff from agricultural and urban land uses. There is a variation in temperature in a given river through they are similar region, may vary with location of sampling sites. The minimum mean temperature determined at upstream (22.6°C) sites might be explained by the highest upland plantation cast canopy cover (90%), high altitude and measurement in the late afternoon. This may slightly lower the temperature. In this regard, the highest temperature recorded in midstream sites (29.3°C) this might be due to lack of canopy cover (0%); the sites have open waters directly exposed to the solar energy, which can attribute to the increase temperature. All the temperature values found to meet the ambient standard (5-30°C) for surface water of Ethiopia.

Small amount of chloride are essential plants and animal life in combinations with a metal such as sodium (Abdullah-al et al, 2018). But, it is detrimental toxic effect above the required limit. The mean concentration chloride measured at upstream, midstream and downstream is 6, 28.9 and 22.6 respectively.

As reported by (Ii, Whiles, Illinois, & David, 2007) anthropogenic impacts and urban activities have long been negative affect aquatic habitat this is true for this study. The dramatic impacts that humans have had on water ecosystem is exemplified in the midstream, the degree of which varied on scale. All most all midstream and some downstream have gotten poor habitat might be due to under pressure from poor waste management from the town administration such as untreated waste discharge, and intensive anthropogenic practices like deforestation, sand and stone dredging, vegetation clearance, grazing and river bank trampling were the most common cause of catchment degradations of the river.

The relatively higher load of pollution correlated closely with decreased pollution sensitive species diversity (like EPT) and increased abundance of number certain pollution tolerant macroinvertebrates like Chironomidae, Physidae, Planorbida, in the midstream and some downstream sites of Lahi River. The result of the current study were quite concurrent with a few earlier studies (Berger et al, 2018;Arimoro & Muller, 2010), which reported that diversity or richness of tolerant species increased and sensitive species dominance decreased considerably a result of anthropogenic threats such as waste discharge, unplanned urbanization, sand and stone derange and other interrelated activity. Land-use patterns and degree of urbanizations influence species composition of river (Luo et al., 2017). Many taxa including Gompidae, Aeshnidae, Coenagrionidae, perlidae, Gyrinidae, Caenidae, Notoectidae, Heptagenidae, Hydrophilidae, Hydropsychidae, Libellulidae, Naucoridae, Elmidae, Athericidae, Chloroperlidae, Calopterygidae, Ceratopogonidae, Corixidae and Psychomyiidae were collected exclusively at midstream sites where as low in numbers at downstream sites. This might be attributed to better physical habitat (bank stability, vegetations protection and substrate condition. It has been reported benthic macroinvertabrates are vulnerable change to habitat conditions (Yi et al., 2018). Taxa richness increase with increasing habitat diversity, suitability, water quality and ecological integrity (Oi, Atano, Egishi, & Anada, 2013). This might also be attributed to differences in ecoregions comprising the upstream, midstream and downstream sites.

Pollutions are not the only cause for disappearance of macroinvertabrates, but physical habitat quality such as substrate composition, vegetation protections and turbulences (Shi et al., 2017). Thus, the poor physical habitat (Table 3) and water quality or ecological integrity condition might be responsible for failure in collected benthic macroinvertabrates taxa from all midstream as well as S9 and S10 (i.e. from downstream sites). The lowest taxa richness noted at midstream might be attributed waste water discharged from the cloth washing and open bathing, municipal waste disposal and organic enrichment through surface run off from urban land use. The highest taxa richness at S1 and S2 the probable reason might be explained due to the sites have been good physical habitat quality (i.e. substrate composition protected riparian vegetations, bank stability, vegetation and canopy cover) water quality as well as good ecological integrity. Abundance of EPT taxa (*perlidae, Batidae, Heptagenidae, Caenidae, Hydropsychidae, Chloroperlidae and Psychomyiidae)* showed that in upstream sites but, disappeared in midstream sites except *Batidae*; this might indicate to decline physical habitat and water quality. However, *Perlidae, Canada, Chloroperlidae* and *Psychomyiidae* were obtained only in upstream sites where as *Batidae, Hydropsychidae* and, *Heptagenidae* was also recorded in downstream sites at S11.*Batidae* was common to all in the three stream whereas, Hydropsychidae, *Heptagenidae* and *Batidae* were the common EPT taxa in upstream and downstream sites. *Canada* was confined to S1 and S2 associated with vegetations and found to be more sensitive to water quality compared to Batidae. A previous study investigating the relationship between macroinvertebrates and environmental factors by (Mereta, 2013) suggested that *Canada* were highly correlated with vegetation and mainly found at sites with good water quality.

FBI value of 3.7 is in the range of 3.76-4.25 which indicated that the water quality of upstream is very good with possible slight organic pollution and downstream (FBI = 5.7) is the range of 5.01-5.75 indicating that the water quality is moderate with fairly substantial pollution where Midstream (FBI=6.8) is in the range between 6.51-7.25 indicating poor river water quality and very substantial degree of pollution (Hilsenhoff, 1988). The index positively correlated BOD and nitrate.

Upstream BMWP value of 73.6 is in the range of 71-100 which indicated that good ecological water quality. Midstream (BMWP=20.6) is in between the BMWP range of 11-40 indicated poor or degraded ecology of river and significant degree of organic pollution. However, downstream (BMWP =43.3) in the range of 41-70 which indicated moderately impacted (Walley and Hawkes, 1997). The score negatively correlated with Nitrate, BOD and Phosphate.

In general, many of the metrics/indices like taxa richness, BMWP, FBI, EPT richness, %EPT, %Deptera and CLI were used in order to estimate the level of ecological disturbance which indicated that at midstream sites is poorly polluted water ecology, however at downstream moderate impacted whereas at upstream is good ecological integrity. The macroinvertabrates community structure and function change naturally along a river according to the stream order, which controls food supply, light and temperature as pointed out in the river continuum concept (Statzner & Higler, 1985). Nevertheless, the changes observed at midstream sites, in the present study at Lahi River may not be natural, because of the total disappearance of the sensitive taxa and some other taxa and, the dominance of the few more tolerant taxa at reach mainly S5, S7, S8 S10.

Cluster analysis using macroinvertabrates data groped S1 with S2 and S7 with S8 (Fig.9) indicating the similarity in water quality or ecological integrity between the paired sites. The analysis based on physicochemical data placed S1, S2 with S3 and S7, S8 with S10 in the same cluster indicating the similarity in water quality among the paired sites. Thus, the cluster analyses output based on macroinvertabrate and physicochemical data indicated the similarity between the two upstream sites S1 and S2, the two midstream sites S7 and S8 in terms of water quality. The analysis farther indicated differences in ecological integrity or water quality among the rest sampling sites and hence generally placed sites according to their pollutions gradient.

None metric multidimensional scaling plots using macroinvertebrates data. Three major groupings, where sites S1, S2, S3, S4 and S11 are with better assemblage of benthic macroinvertabrates which might attributed to low organic matter lode due to low number of human activity and husbandry house therefore, better ecological integrity. However in S5 and S6 are impacted sites, and while in S7, S8 and S10 are those receiving point source discharges from the catchments.

The variances of macroinvertabrates taxa explained by the first CCA axes exceeded. CCA tri-plot of macroinvertabrates data clearly separated the midstream sites with disturbances water quality or ecology i.e. S7 and S8. The trip-lots for macroinvertabrates assemblage separated S5 and S6 from other sampling sites. The direction proportional influences of EC, BOD, turbidity, chloride and TDS was pointing towards the midstream and some downstream i.e. S10 the figures farther revealed that the direction proportional influence of DO pointing towards upstream. The direction of the most physicochemical parameters in the tri-plot were away from S1, S2, S3 as well as S11since the measured values of those variables that increase with increase in pollutions load were lower at these sites

Assessment of ecological status of Lahi River using physicochemical parameters, physical habitat, biotic indices/metrics, human activity and multivariate analyses indicated ecological deterioration seen poorly at midstream and moderately at downstream, where very good ecological integrity at upstream sites. Untreated or poorly treated waste effluent discharge, institutions, commercial center, crop productions as well as sand and stone dredging, cloth washing, open bathing, and waste water discharge from the town land uses all along the length of the river were the major environment stresses that affect ecological integrity of Lahi River. This is in agreement with reports from studies conducted in other rivers in the county (Desalegne, 2018; Hailu and Legesse, 1997; Legesse,200).

## Conclusions and Recommendations

### Conclusion

We live in a world of varied ecosystems and our existences would not be possible without the life-supporting services one another. Indeed, rivers are providing a range of biological, environmental and socio-economic benefits for indigenous population. The result of physicochemical, macroinvertabrate and habitat assessment of this river from different sampling stations indicated that the higher level of BOD and lower DO occur at the midstream where much influence of Finoteselam town town, like solid and liquid wastes disposal, sand and stone dredging, poor farming method, distraction of riparian forest, car and cloth washing, open bathing and poor solid and liquid waste municipal management and sewage discharge were the major environmental stressors responsible for ecological deterioration. This correlates with higher values of pollution indicating nutrients (nitrate and phosphate). Generally this study shows that midstream was relatively more polluted due to discharge of solid and liquid wastes from the town and intensive agricultural activity. The influence of Finoteselam town mentions above has directly caused the considerable reduction of macroinvertabrate diversity.

### Recommendations

- Reducing directly piped drainage connection using infiltration and retention as well as proper solid waste management might be a logical step in the mitigation of the impacts of Finoteselam on Lahi River.
- Municipality of finoteselam town should be preparing solid and liquid wastes disposal site.
- Attempts to improve ecological status of the Lahi River should include regulation of point source loading such as waste water discharge from olid and liquid waste from the town and effective water shaded management which includes riparian forest improvement to stop non-point source loading.
- Schistosomiasis assessment of LahiRiver water should be working out as children, women and local community were found to heavily depend on the river water for taking bath, swimming and washing at different sites including the polluted ones

## Ethical Considerations

Ethical clearance was obtained from BUERB

## Availability of data

Supporting data for the current study are available from the corresponding author on reasonable request.

